# Preclinical evaluation of ixazomib for high-risk pediatric brain tumors

**DOI:** 10.64898/2026.09.11.750478

**Authors:** Elizabeth Janko, Faye M. Walker, Mathew Slade, Etienne Danis, Pradeep Bompada, Lays Martin Sobral, Julia Chapman, Ishmeen Kaur, Simran Bal, Gabrielle Link, Nicholas McQuillan, Jean M. Mulcahy Levy, Adam L. Green, Nathan A. Dahl

**Affiliations:** Morgan Adams Foundation Pediatric Brain Tumor Research Program, Department of Pediatrics, University of Colorado School of Medicine, Aurora, CO; Biostatistics and Bioinformatics, University of Colorado School of Medicine, Aurora, CO; Center for Cancer and Blood Disorders, Children’s Hospital Colorado, Aurora, CO

**Keywords:** Ixazomib, proteasome inhibition, medulloblastoma, atypical teratoid/rhabdoid tumor, diffuse midline glioma

## Abstract

**Background:** Many of the highest risk pediatric brain cancers continue to experience poor clinical outcomes despite intensification of current multimodal therapy. Proteasome inhibition has shown preclinical promise across a range of cancer models including diffuse midline glioma (DMG), medulloblastoma, and atypical teratoid / rhabdoid tumors (ATRT), though clinically viable agents have been limited. Recent studies in adults with glioblastoma suggest that ixazomib, a second-generation proteasome inhibitor, might achieve therapeutic concentrations in the CNS, presenting the opportunity that a CNS penetrant proteasome inhibitor might be similarly leveraged for benefit in childhood brain cancers.

**Methods:** Ixazomib was tested against cell lines and orthotopic xenograft models of DMG, Myc-amplified medulloblastoma (Myc-MB), and ATRT. RNA sequencing and LC-MS based proteomics were utilized to define functional consequences of ixazomib treatment in these models. Proteasome activity readouts were used to assess ixazomib activity across brain regions and extracranial solid organs.

**Results:** Ixazomib demonstrates consistent cytotoxic effect across high-risk brain tumor models at low nanomolar concentrations. Ixazomib treatment activates proteostatic stress response and apoptosis. Treatment with ixazomib does not demonstrate survival benefit in orthotopic models, however, and pharmacodynamic testing suggests insufficient inhibition of proteasome activity within the CNS compared to extracranial tissues.

**Conclusions:** While many pediatric brain tumor models demonstrate susceptibility to proteasome inhibition, ixazomib may lack sufficient blood-brain barrier penetration to be a translationally viable means of exploiting this vulnerability.

**Key Points:**

- Ixazomib is effective against Myc-MB, ATRT and DMG tumors *in vitro*
- Ixazomib treatment does not prolong survival in orthotopic models
- Ixazomib may fail to achieve sufficient suppression of proteasome activity within the CNS

**Importance of Study:** Many high-risk subsets of pediatric brain tumors continue to experience unsatisfactory clinical outcomes under current treatment regimens. Previous research has shown many of these diseases to be responsive to proteasome inhibition, and recent phase 0 data in glioblastoma has suggested that the agent ixazomib may achieve meaningful concentrations in the CNS following systemic administration. In this study, we test ixazomib across a range of pediatric brain tumor models. Using transcriptomic and proteomic approaches, we find that ixazomib effectively triggers proteostatic stress responses and induces apoptosis across a histologic range of CNS malignancies. Orthotopic models fail to demonstrate a survival benefit from ixazomib treatment, however, and pharmacodynamic readouts suggest limited proteasome suppression within the CNS. These findings caution further translational efforts for this agent in children with primary brain tumors.

## Introduction

Considerable progress has been made in recent years for many pediatric brain tumors, though improvements in clinical outcomes have not been uniformly distributed. Patients with high-risk subsets of disease continue to experience unsatisfactory response rates to current therapies, and development of novel therapeutics for these malignancies remains an unmet need.

The ubiquitin-proteasome system (UPS) is responsible for the controlled degradation of intracellular proteins in service of maintaining protein homeostasis. Poly-ubiquitination of proteins by E3 ligases targets them to the 26S proteasome complex, allowing for directed turnover into smaller peptide fragments^1–3^. Dysregulation of the UPS has been implicated in tumorigenesis, both in altering the balance of oncogenes versus tumor suppressor proteins as well as through prevention of apoptosis normally triggered by prolonged proteostatic stress^3,4^. This has led to considerable interest in proteasome inhibition as a therapeutic strategy, which has been extensively reviewed elsewhere^1,2,5^. For primary brain tumors, exploitation of proteasome dependence has been constrained by the limited penetration of most clinical grade agents across the blood-brain barrier (BBB)^6^. The exception to this is marizomib, a second-generation proteasome inhibitor (PI) with a more lipophilic structure, facilitating confirmed CNS distribution in rats and non-human primates^7^. Despite promising preclinical signal across a range of pediatric and adult primary brain tumors, however, a phase 3 study in adults with newly diagnosed glioblastoma failed to demonstrate a survival benefit with the addition of marizomib to the standard backbone of radiation plus temozolomide^8^. Clinical development of this agent has now stalled, leaving the future of proteasome inhibitors for primary brain tumors uncertain^9^.

Ixazomib (MLN2238) is another second-generation proteasome inhibitor, comparatively unique for its bioavailability following oral administration^5^. Like other non-marizomib PIs, ixazomib was not predicted to readily cross the BBB nor extensively tested against primary CNS malignancies. However, a recent phase 0 study in adults with glioblastoma (NCT02630030) found measurable ixazomib concentrations within tumor tissue at time of surgical resection^10^, supporting the development of phase 1/2 clinical trials for this indication^11^. The potential therapeutic utility of ixazomib for high-risk pediatric brain tumors remains unexplored.

In this study, we test the efficacy of ixazomib against models of diffuse midline glioma (DMG), Myc-amplified medulloblastoma (Myc-MB), and atypical teratoid / rhabdoid tumor (ATRT), three pediatric CNS malignancies that have previously been shown to be PI-responsive^6,12,13^. Utilizing both cell lines and representative orthotopic xenograft models, we investigate mechanisms of anti-tumor effect and probe efficiency of ixazomib penetration across the BBB, all with the goal of determining whether ixazomib warrants further translational study in children diagnosed with PI-responsive CNS malignancies.

## Materials and Methods

### Western Blotting

Western blotting was performed as previously described^14^. Antibodies used were as follows, prepared according to manufacturer recommendations: Cleaved Caspase-3 (Asp175) Rabbit mAb Cell Signaling #9661, PARP (46D11) Rabbit mAb Cell Signaling #9532, Ubiquitin (P37) Rabbit Cell Signaling #58395, p21 (12D1) Rabbit mAb Cell Signaling #2947), β-Actin (13E5) Rabbit mAb Cell Signaling #4970.

### Orthotopic Xenograft Models

Female athymic nude mice aged 4 to 8 weeks were anesthetized, immobilized and prepared for intracranial injection as we have previously described^15,16^. All cell lines were prepared in 2 μL serum-free RPMI for each injection. To target the cerebellum, a 1.0 mm diameter burr was drilled in the cranium using a Dremel drill outfitted with a dental drill bit at 1.500 mm to the right and 2.000 mm posterior to lambda, with cell suspension injected 3.000 mm ventral to the surface of the skull. To target the pons, a 1.0 mm diameter burr was drilled in the cranium at 1.000 mm to the right and 0.800 mm posterior to lambda, with cell suspension injected 5.000 mm ventral to the surface of the skull. The burr hole was sealed, incision closed, and topical antibiotic. Post-surgical pain was controlled with 5 mg/kg/day SQ carprofen on the day of and for 2 days following the procedure. Protocol-defined endpoints included ataxia, >15% weight loss, or significant distress. University of Colorado Institutional Animal Care and Use Committee (IACUC) approval was obtained and maintained throughout the conduct of the study.

### Combination Indices

Cells were seeded in suspension conditions in flat-bottom, ultra-low attachment 96 well plates (Corning) at approximately 20,000 cells per well in 90 µL of cell culture medium. The same day, cells were treated in triplicate with 10 µL of DMSO or 4-9 drug concentrations in fixed-ratio by serial dilution, and/or fixed-dose X-ray irradiation, determined by monotherapy data such that doses within an order of magnitude of the combined null (theoretical model indicating neither synergism nor antagonism) IC50 would appear towards the center of the dose-response curve. Cells viability was determined at five days using MTS [3-(4, 5-dimethylthiazol-2-yl)-5-(3-carboxymethoxyphenyl)-2-(4-sulfophenyl)-2H-tetrazolium] (CellTiter 96 AQueous One Solution [Promega, Madison, WI, USA]). Dose-response data were first entered into CompuSyn (Chou, 2005, ComboSyn Inc.) to obtain Combination Index (CI) values^17^. Then, the synergy^18^ package was used to recover CI values for validation and batch processing in Python. Synergy was determined on a dose-dependent basis based on the CI value. CI plots were visualized first in CompuSyn and then Python (Seaborn^19^, Matplotlib^20^, NumPy^21^, and synergy^18^ packages).

### RNA sequencing and Proteomics

RNA library construction, sequencing, and downstream analyses were performed as we have previously described^22^.

For proteomic analysis, cells were washed 3x with cold PBS and frozen in Eppendorf MS-compatible microcentrifuge tubes. LC-MS proteomics was performed by the CU Anschutz Mass Spectrometry Proteomics Shared Resource (RRID: SCR_021988) on the Bruker timsTOF Pro platform. Proteomics data analysis was performed using the DEP2 package (v1.8.0)^23^. Protein intensities were log2 transformed and normalized using Variance Stabilizing Normalization (VSN). To account for missing values, the Bayesian Principal Component Analysis (BPCA) imputation method was applied. Differential expression analysis was conducted by comparing Treated and Control groups using the manual linear model test provided by DEP2. P-values were adjusted for multiple testing using the Benjamini-Hochberg (BH) procedure. Significantly differentially expressed proteins (DEPs) were defined based on an adjusted p-value < 0.05 and an absolute fold change (FC) > 1.25.

Over-Representation Analysis (ORA) was performed using clusterProfiler (v4.14.0)^24^ against gene sets from the msigdbr package (v7.5.1)(https://cran.r-project.org/web/packages/msigdbr/), including Reactome, WikiPathways, KEGG, and GO:BP collections. To visualize the relationship between statistical significance and biological function, a customized volcano plot was generated using the EnhancedVolcano package (v1.24.0). Specific proteins belonging to key enriched pathways, including Proteasome inhibition (GO:BP), Cell Cycle (Reactome), TP53 Transcriptional Regulation (Reactome), and Translation (Reactome) were color-coded and labeled to highlight their distribution across the differential abundance profile.

Additional visualizations, including hierarchical clustering heatmaps and gene-concept networks (cnetplots), were produced using pheatmap (v1.0.12) and enrichplot (v1.26.0), respectively. All data manipulation and figure formatting were carried out using dplyr (v1.1.4) and ggplot2 (v3.5.1).

### Chymotrypsin-Like Activity Assay

Non-tumor-bearing athymic nude mice were administered ixazomib 20 mg/kg or vehicle control. After 1 or 12 hours, mice were euthanized and cortex, brain stem, and cerebellum regions were harvested. Tissues were cut into small pieces with a razor blade in RPMI serum-free media. Tissue was then passed through 50 micron cell strainers into 50 mL conical tubes (Falcon), tissue solutions were centrifuged at 1,500 RPM for 5 minutes, and supernatant was aspirated. Single cells were resuspended in RIPA buffer. Protein quantities were standardized through BCA protein quantification assay. Using a 96-well plate, each sample was tested in triplicate using the chymotrypsin-like Proteasome-Glo Assay (Promega Cat No G8622). Absorbance was read at 492 nM on a BioTek Synergy H1 microplate reader (BioTex Instruments, VT).

## Results

### Ixazomib is effective against high-risk pediatric brain tumors *in vitro*

In order to begin assessing the potential for ixazomib as a viable therapeutic option for pediatric brain tumors, we first tested it against cell culture models of DMG, Myc-MB, and ATRT. Dose response curves demonstrated mean inhibitory values at 72 hours consistently in the low nanomolar range against all models tested. Mean inhibitory values for DMG lines ranged from 10 to 32 nM (10 nM for SU-DIPG-IV, 26 nM for HSJD-DIPG-007, and 32 nM for BT245) (**Figure 1A**). Myc-amplified medulloblastoma lines exhibited response at a range from 12 nM to 46 nM (12 nM for D283, 46 nM for D425, and 16 nM for D458) (**Figure 1B**), and ATRT lines responded from 23 nM to 60 nM (31 nM for BT12, 60 nM for BT16, and 23 nM for MAF-737) (**Figure 1C**). These suggest broad responsiveness to ixazomib across tumor types at concentrations that may be clinically achievable.

**Figure 1.**
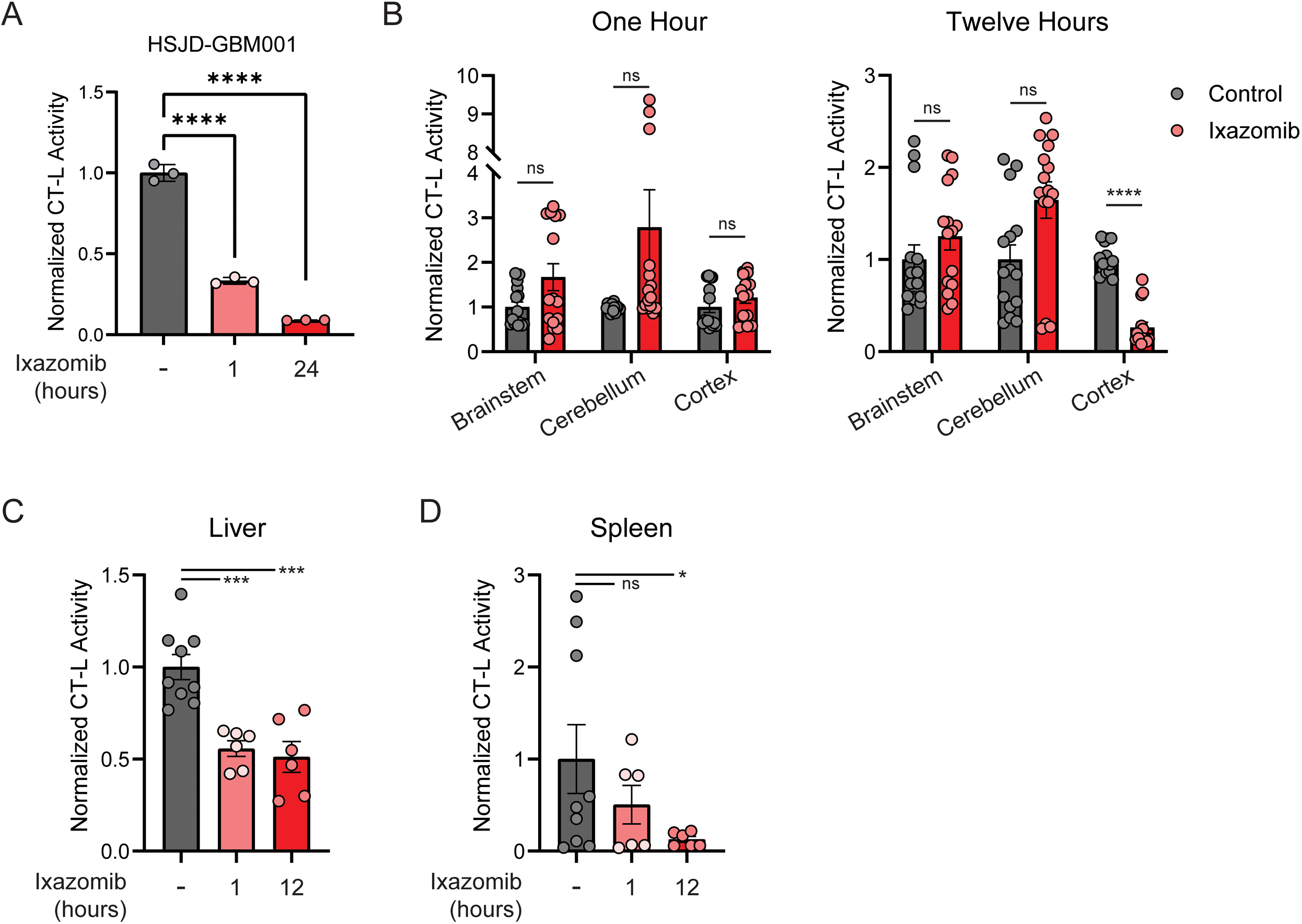
Ixazomib is effective against high-risk pediatric brain tumors *in vitro*. **A.** Mean inhibitory concentrations of ixazomib in diffuse midline glioma cells (SU-DIPG4 10 nM, HSJD-DIPG-007 26 nM, BT245 32 nM). **B.** Mean inhibitory concentrations of ixazomib in Myc-amplified medulloblastoma cells (D283 12 nM, D425 46 nM, D458 16 nM). **C.** Mean inhibitory concentrations of ixazomib in atypical teratoid / rhabdoid tumor cells (BT12 31 nM, BT16 60 nM, 23 nM MAF-737). **D.** Immunoblot for total ubiquitin, cleaved caspase, and cleaved PARP at indicated duration of ixazomib treatment in D458 cells. **E.** Immunoblot for total ubiquitin, cleaved PARP, and p21 following ixazomib 100 nM 24 hour treatment in indicated cell lines.

Inhibition of proteasome function should impair clearance of polyubiquitinated proteins and ultimately lead to induction of apoptosis. To test the kinetics of this, we performed immunoblotting for total ubiquitinated protein in D458 cells at timepoints ranging from 0 to 24 hours following 100 nM ixazomib exposure. We observed a progressive accumulation of ubiquitinated protein over this time course, with cleavage of the apoptosis-associated proteins caspase 3 and PARP occurring between 16 and 24 hours (**Figure 1D**). We then repeated these assessments following 24 hours of treatment in additional DMG, ATRT, and Myc-MB cell lines, finding similar evidence of ubiquitinated protein accumulation, induction of apoptosis (cleaved PARP), and cell cycle arrest (p21) (**Figure 1E**). Taken together, these data support ixazomib as a potent agent for triggering cell death across a histologic range of high-risk pediatric brain tumors.

### Ixazomib treatment is synergistic with radiation therapy

Despite well characterized late effects in children, ionizing radiation remains a mainstay of therapy for high-risk pediatric brain tumors. For some tumors such as DMG, radiation therapy (RT) is the only broadly accepted standard of care. In light of this, we assessed whether treatment with ixazomib would elicit synergistic, additive, or antagonistic anti-tumor effects when combined with RT. We exposed a panel of DMG culture models to increasing concentrations of ixazomib in the presence or absence of a fixed RT dose, determined according to prior cell line-specific RT responses (range 4 Gy – 10 Gy). Combination indices were generated by analyzing dose response data according to the Chou-Talalay model^17^. This demonstrated a significant synergistic effect across 9 of the 11 DMG models tested (**Figure 2A**), supporting the use of PI in combination with radiation therapy.

**Figure 2.**
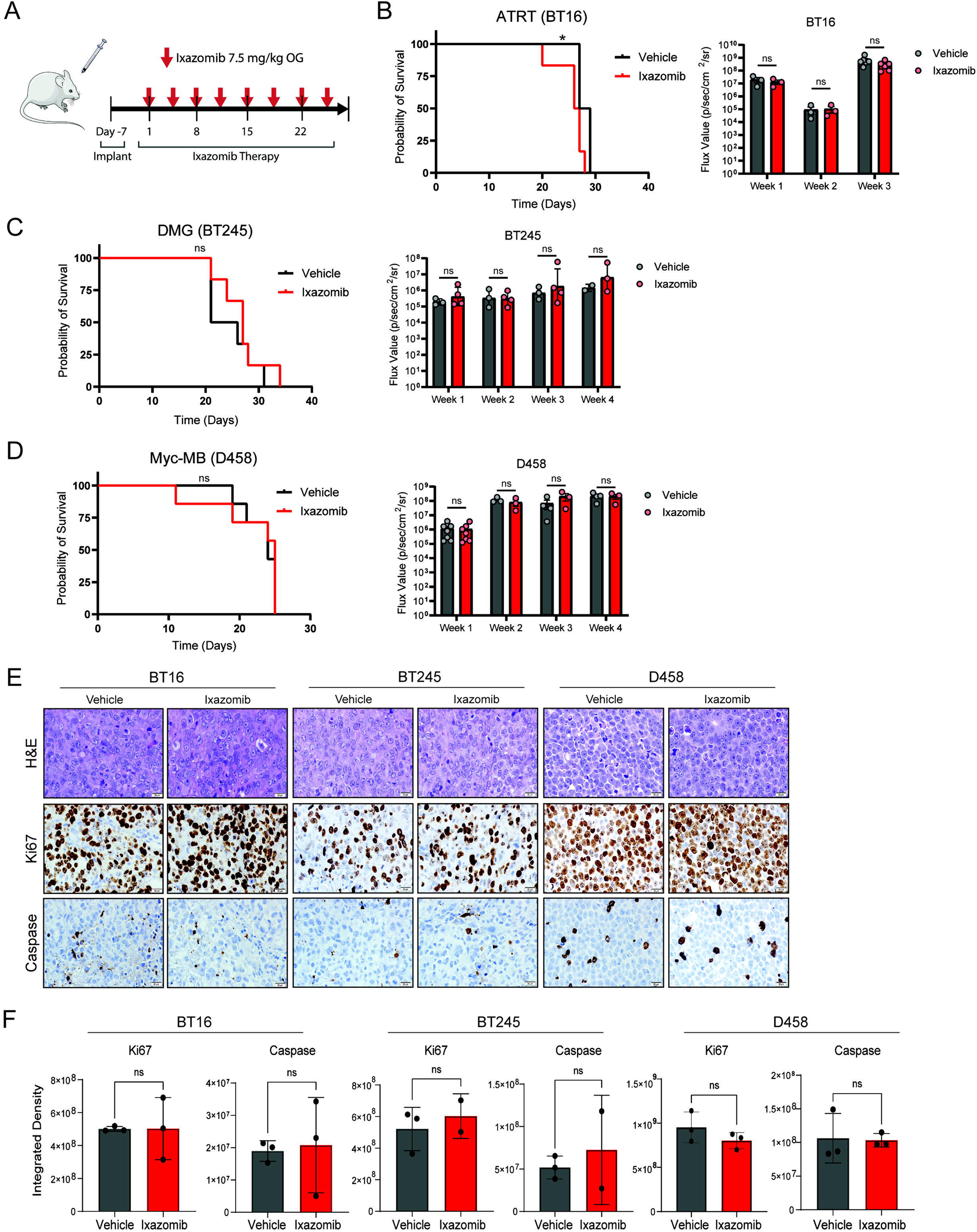
Ixazomib is synergistic with radiotherapy. **A.** Combination Index (CI) values for ixazomib with radiation across indicated pHGG cell lines. CI values between 0 and 1 are synergistic; the closer to 0, the more synergistic. CI values of 1 are additive, and CI values above 1 are antagonistic **B.** Colony formation (left) and quantification (right) of SU-DIPG4 cells treated with DMOS, IC25, or IC50 dose of ixazomib at indicated radiation doses. **C.** Colony formation (left) and quantification (right) of HSJD-DIPG007 cells treated with DMOS, IC25, or IC50 dose of ixazomib at indicated radiation doses. Comparisons in (B) and (C) reflect pairwise two-tailed Student’s t-tests (ns p=>0.05, * p=<0.05, ** p=<0.01, **** p=<0.000).

To further test the potential therapeutic utility of combining ixazomib with radiotherapy, we performed clonogenic assays at variable ixazomib and radiation doses in two DMG cell lines, SU-DIPG4 and HSJD-DIPG007. In each cell model, we observed a dose-dependent decrease in colony formation from either treatment alone, with the greatest decrease seen when treatment modalities were combined (**Figure 2B-C**). Taken together, these data suggest that ixazomib might provide benefit to current treatment backbones incorporating the use of radiation therapy.

### Ixazomib treatment disrupts common oncogenic programs across brain tumor models

Given the potent anti-tumor effect we observed from ixazomib across a range of brain tumor models, we next sought to characterize the transcriptional and post-transcriptional consequences of ixazomib treatment for these cancers. We took cultures of HSJD-DIPG-007 (DMG), D425 (Myc-MB), and BT12 (ATRT) and treated with ixazomib 250 nM for 4 hours before performing RNA sequencing (n=3 each). This identified 2312 to 3880 genes differentially expressed following ixazomib treatment (3880 HSJD-DIPG-007, 3799 D425, 2312 BT12) (LFC ≥1.5, p adj <0.05) (**Figure 3A**). Gene set enrichment analysis performed individually for each cell model showed downregulation of programs involved in cell cycle progression and upregulation in processes governing apoptosis and protein catabolism in response to stress (**Figure 3B** and **Supplementary Table 1A**). Many identical or substantially similar gene sets were significantly enriched across multiple tumor models, so we performed an integrative analysis to identify conserved ixazomib effects. Using the Metascape cross-dataset pipeline^25^, we found marked overlap between common genes and enriched biological processes up-or downregulated across tumor types (**Figure 3C** and **Supplementary Table 1B**). Commonly upregulated gene ontology (GO) terms included immune activation (e.g. cytokine and TNFα-NFκB signaling, hallmark inflammatory response), stress response (KEAP1-NFE2L2 pathway, response to external stimulus), p53 activity, and apoptosis. Commonly downregulated GO terms predominantly included processes related to cell cycle progression (e.g. microtubule binding, hallmark E2F targets, regulation of chromosome separation) and glioneuronal differentiation (modulation of synaptic transmission, cell morphogenesis, neuronal system) (**Figure 3D-E** and **Supplementary Table 1C-D**). We then turned to the transcription factors (TFs) likely responsible for organizing these downstream transcriptional changes. Utilizing the transcriptional regulatory relationships unraveled by sentence-based text-mining (TRRUST) database^26^, we nominated TFs from the integrated transcriptomic data. These included XBP1 and ATF4, TFs with established roles in unfolded protein and integrated stress response^27–29^, as well as RELA and NFκB, inflammatory signaling modulators with previously defined responsiveness to proteasome inhibition^30,31^ (**Figure 3F** and **Supplementary Table 1E**). Together, these data define core oncogenic programs that are disrupted by ixazomib treatment, agnostic of the diverse underlying tumor biology.

**Figure 3.**
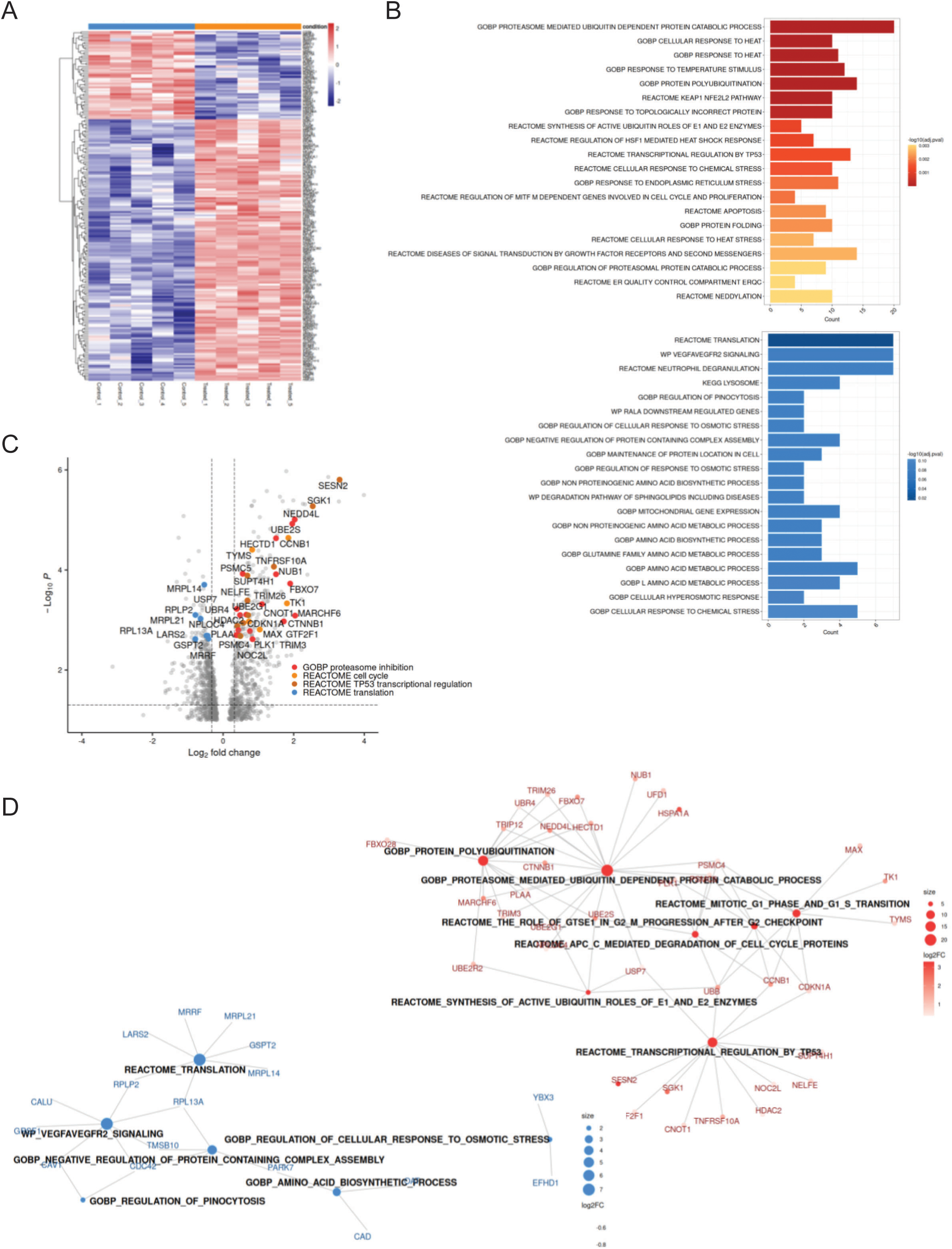
Ixazomib treatment disrupts common oncogenic transcriptional programs. **A**. Unsupervised hierarchical clustering of HSJD-DIPG-007, D425, and BT12 gene expression LFC ≥ 2 in control vs ixazomib-treated samples (n=3 each, p<0.001). **B**. Gene sets enriched (FDR padj <0.05) in (B). Dark colored circles (red=up, blue=down) denote gene sets significantly enriched in three cell models, light colored circles denote gene sets enriched in two cell models, while grey indicates terms unique to one model. **C**. Circlos plot of differentially up-(top) or downregulated (bottom) genes (top 1000 genes, p adj <0.05) in HSJD-DIPG-007, D425, and BT12 models. Purple connections indicate identical genes in both lists, while blue connections indicate common enriched GO terms. **D**. Clustered heatmap of GO terms enriched from (D). **E**. Representative common gene sets from RNA-seq in HSJD-DIPG-007 (left) and D425 (right) models. **F**. TRRUST inference of transcription factor-target pairs from genes differentially downregulated following ixazomib treatment, ranked by Fisher’s exact test (-log(p val)).

While transcriptomic readouts provide insight into the cellular phenotype affected by ixazomib treatment, proteasome inhibition mechanistically works at the post-translational stage. To address this, we augmented these experiments by performing mass spectrometry-based proteomics from the BT12 model following ixazomib treatment (n=6 each). Differential expression analysis utilizing the DEP2 pipeline (see Methods) identified 203 proteins significantly differentially expressed following ixazomib treatment (LFC ≥1.25, p adj <0.05) (**Figure 4A**). Pathway enrichment identified upregulation of programs involving ubiquitin-mediated protein degradation (e.g. proteasome mediated ubiquitin dependent protein catabolism, protein polyubiquitination, protein folding), stress or heat shock response (response to heat, response to temperature stimulus, KEAP1-NFE2L2 pathway, HSF1-mediated heat shock response), regulation by p53, and apoptosis. Downregulated proteins were most strongly enriched in processes related to protein translation, amino acid synthesis, and protein complex assembly (**Figure 4B-D**). These proteomic networks demonstrate that ixazomib treatment in ATRT impairs ubiquitin-mediated protein degradation to drive stress response and ultimately activation of apoptosis.

**Figure 4.**
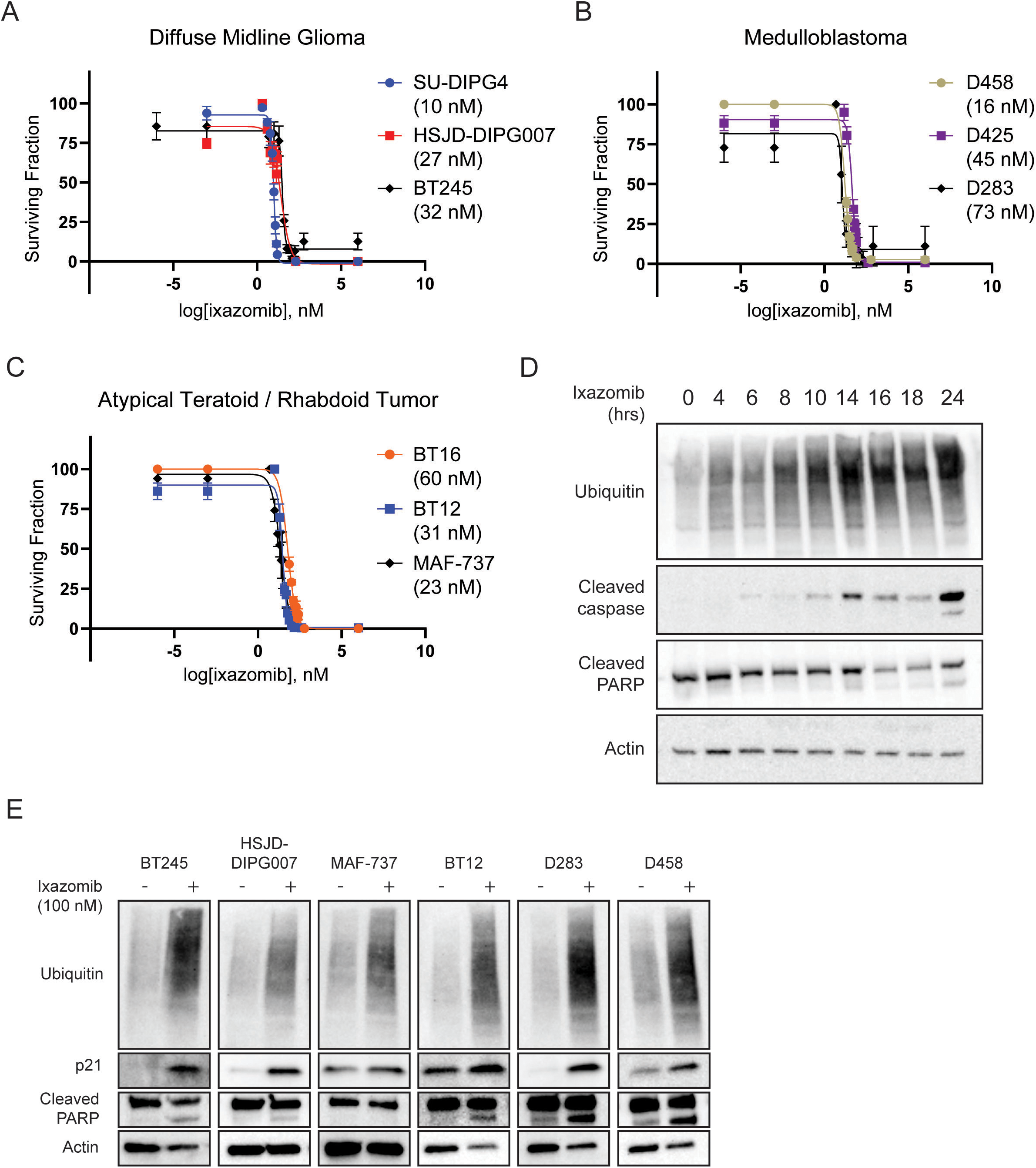
Ixazomib treatment alters proteome homeostasis to drive stress response and apoptosis. **A**. Unsupervised hierarchical clustering of LC-MS proteomics from BT12 LFC ≥1.25 in control vs ixazomib-treated samples (n=5 each, p<0.05). **B**. Gene ontology enrichment of differentially expressed proteins from (A). Upregulated processes are represented in red (top), while downregulated processes are represented in blue (bottom). **C**. Volcano plot of differential protein expression following ixazomib treatment. Proteins associated with proteasome inhibition, cell cycle regulation, regulation by TP53, or protein translation are highlighted. **D**. Gene ontology network constructed from differentially expressed proteins. Upregulated programs are in red, downregulated programs in blue. Nodes denote enriched processes, while connections indicate specific differentially expressed proteins.

### Ixazomib treatment does not provide survival benefit *in vivo*

Given the promising anti-tumor effects of ixazomib treatment *in vitro*, we next tested whether these findings would translate to a survival benefit in orthotopic xenograft models. Ixazomib has been previously tested in murine models at a range of doses using both oral and tail vein administration, though the clinical product is FDA approved for oral use only. We found the published^32^ regimen of 7.5 mg/kg by oral gavage (OG) twice weekly to be tolerable in tumor-bearing athymic nude mice, while doses at 10 mg/kg or above elicited protocol-defined dose limiting toxicity (not shown). We then generated orthotopic models by injecting D458 medulloblastoma cells or BT16 ATRT cells into the cerebellum or BT245 DMG cells into the pons before randomization to ixazomib treatment or vehicle control (**Figure 5A**). Unfortunately, mice treated with ixazomib did not demonstrate a significant survival benefit or decrease in tumor size by BLI in any model system tested (**Figure 5B-D**). To assess evidence of activity within the target tumors, tumor tissue was harvested and subjected to histology processing and immunohistochemistry staining. We observed no qualitative difference in tumor histology nor significant difference in quantification of Ki67 or caspase 3 activation when comparing ixazomib treated samples to controls within a tumor group (**Figure 5E**).

**Figure 5.**
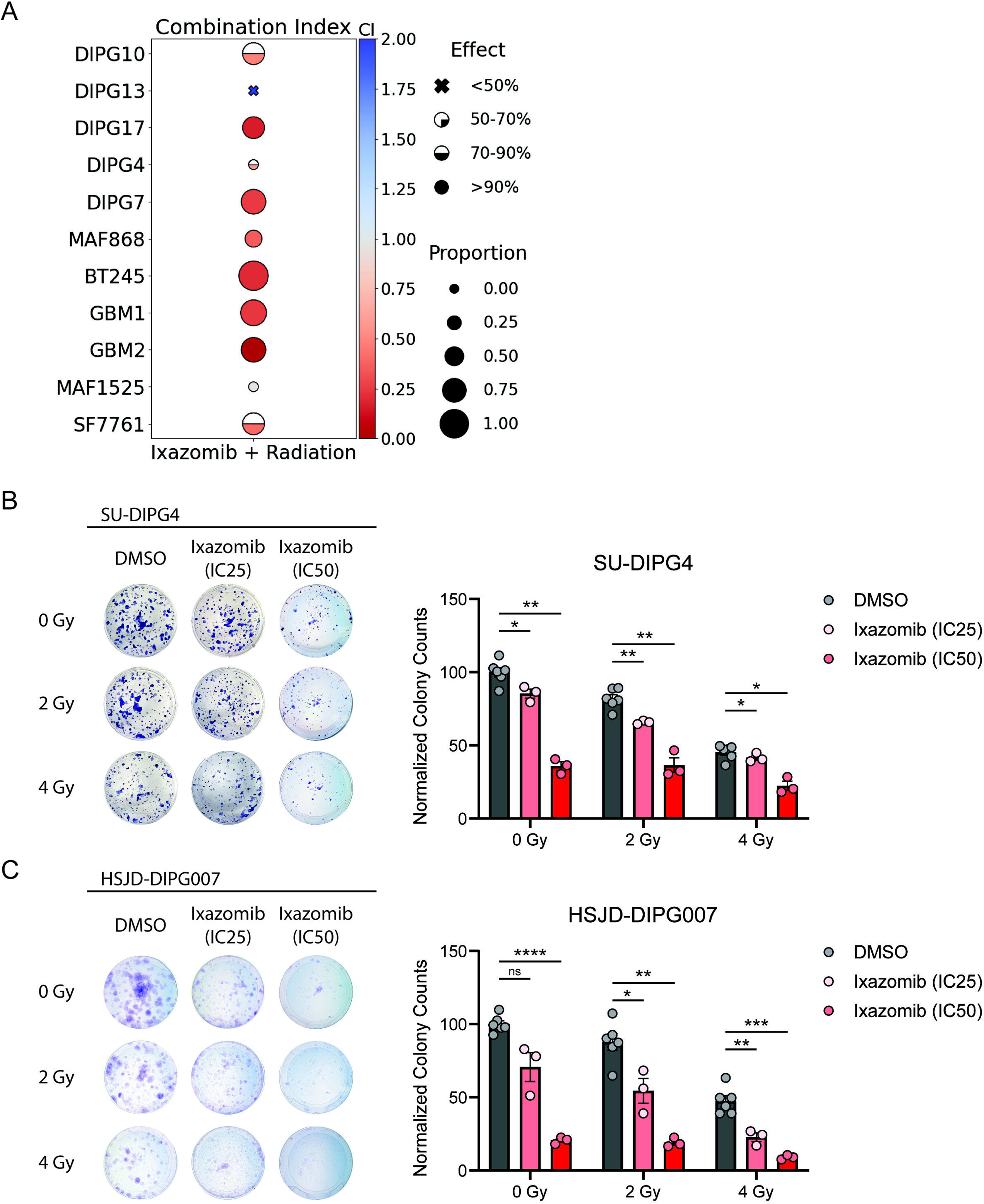
Ixazomib treatment does not prolong survival in orthotopic models of pediatric brain tumors. **A.** Schema of treatment delivery for ixazomib in intracranial xenograft models. **B.** Kaplan-Meier survival analysis (left) and weekly BLI values (right) of intracranial BT16 xenografts treated with ixazomib or vehicle control. Comparisons reflect pairwise two-tailed Student’s t-tests (ns p >0.05). **C.** Kaplan-Meier survival analysis (left) and weekly BLI values (right) of intracranial BT245 xenografts treated with ixazomib or vehicle control. Comparisons reflect pairwise two-tailed Student’s t-tests (ns p >0.05). **D.** Kaplan-Meier survival analysis (left) and weekly BLI values (right) of intracranial D458 xenografts treated with ixazomib or vehicle control. Comparisons reflect pairwise two-tailed Student’s t-tests (ns p >0.05). **E.** H&E staining and immunohistochemistry from (B-D) showing Ki67 and cleaved caspase 3. **F.** Quantification from (E). Comparisons reflect pairwise two-tailed Student’s t-tests (ns p >0.05).

We next reasoned that the discordance observed between *in vitro* testing and the orthotopic models may suggest insufficient exposure of ixazomib across the blood-brain barrier to achieve proteasome suppression within target brain regions. To test this, we utilized a pharmacodynamic readout of chymotrypsin-like (CT-L) activity, one of the major proteolytic reactions catalyzed by the proteasome 20S core^33^. This approach has previously demonstrated robust suppression of CT-L activity in the brain one hour following administration of marizomib, a PI with comparatively robust diffusion into the CNS^12^. We first validated the assay with ixazomib *in vitro*, observing clear evidence of CT-L activity suppression at 1 and 24 hours following ixazomib treatment (**Figure 6A**). We then administered ixazomib at a single supratherapeutic dose of 20 mg/kg to non-tumor-bearing mice before euthanizing at 1 and 12 hours, followed by single-cell disaggregation from brainstem, cerebellum, and cerebral cortex. While we noted some signal of suppression within the cortex after 12 hours, we otherwise did not observe consistent evidence of CT-L activity suppression across brain regions or timepoints (**Figure 6B**). In contrast, cells collected from liver and spleen at the same timepoints both demonstrated evidence of CT-L activity suppression following ixazomib administration, confirming the ability of systemic ixazomib administration to suppress proteasome function within extracranial solid organs (**Figure 6C**). Taken together these data suggest that even at supratherapeutic doses, systemic administration of ixazomib does not achieve sufficient CNS penetration to effectively suppress proteasome function within relevant brain regions. These pharmacodynamic findings are unfortunately consistent with the observation that ixazomib treatment at a tolerable dosing regimen does not provide a survival benefit in any intracranial orthotopic model tested.

**Figure 6.**
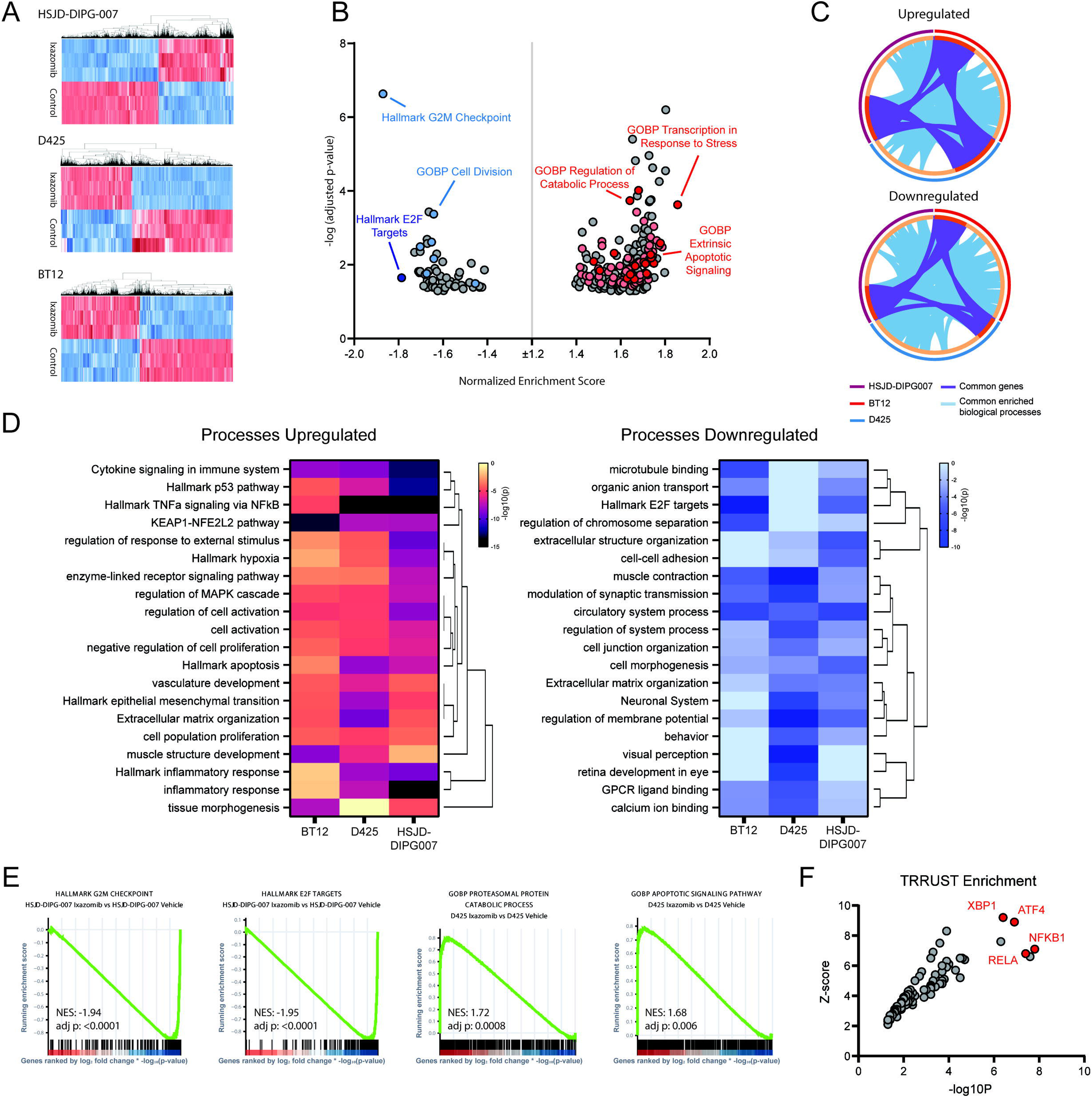
Ixazomib may lack sufficient blood-brain-barrier penetration at tolerated doses. **A.** Chymotrypsin-like (CT-L) activity from HSJD-GBM001 cell lines treated with ixazomib 20 nM at indicated timepoints. **B.** Normalized CT-L activity from dissected brainstem, cerebellum, or cortex at one (left) or twelve (right) hours following ixazomib administration. **C.** Normalized CT-L activity from dissected liver at one or twelve hours following ixazomib administration. **D.** Normalized CT-L activity from dissected spleen at one or twelve hours following ixazomib administration. All comparisons reflect pairwise one-tailed Student’s t-tests (ns p=>0.05, * p=<0.05, ** p=<0.01, **** p=<0.000).

## Discussion

Many high-risk subtypes of pediatric brain tumors are desperately in need of novel therapeutic options. Proteasome inhibition has shown promise across a range of both pediatric and adult brain cancers, but as is often the case for CNS malignancies, the ability of available agents to cross the blood-brain barrier remains a translational limitation. The development of marizomib as the first CNS-penetrant PI led to a flurry of preclinical and early clinical studies within the broader neuro-oncology community despite the agent not ultimately reaching clinical viability^6–9,12,13^. Recent phase 0 testing in adults suggested that systemic administration of ixazomib could achieve measurable concentrations in resected glioblastoma^10^, opening the possibility for parallel clinical application to pediatric CNS tumors previously shown to be PI-responsive^34^.

In this present study, we rigorously test ixazomib across cell and animal models of DMG, Myc-MB, and ATRT, three pediatric cancers that experience unsatisfactory outcomes under current standard of care yet have been shown to respond to proteasome inhibition in previous studies. We find that ixazomib is effective in suppressing proteasome function, activating proteostatic stress responses to drive apoptosis across diverse tumor histology. However, we find no consistent evidence of effective suppression of proteasome activity within the CNS, nor do we observe a survival benefit across three orthotopic xenograft models. These findings are unfortunately consistent with the limited BBB penetration predicted for other second generation proteasome inhibitors^35^. Our data presented is not without limitations. While we utilize biologically relevant pharmacodynamic readouts, we lack robust LC-MS based pharmacokinetic studies of ixazomib distribution in tissues. It is possible that alternative routes of administration or augmented delivery mechanisms such as intracerebroventricular dosing, convection enhanced delivery, or focused ultrasound might increase ixazomib delivery to target brain regions. And it is possible that differences between human and murine BBB^36,37^ might allow for meaningful CNS distribution in patients despite the lack of signal seen here. However, we present these findings as a caution to planned translational efforts of ixazomib for children with primary brain tumors, as a more critical evaluation of CNS distribution may be warranted before advancing to clinical trials in this population.

## Author Contributions

E.J., F.M.W., and I.K. performed the in vitro phenotypic assays. M.S. and G.L. performed the radiation synergy assays. E.J., S.B., and J.C. performed the Western blots. E.J. performed RNA and proteomic sequencing, E.D., P.B., and Pluto Bio performed the bioinformatic analysis. E.J., F.M.W., and N.M. performed the CT-L assays. E.J. and F.M.W. performed the animal experiments. E.J. and N.A.D. prepared the figures and wrote the manuscript. J.M.M.L., A.L.G., and N.A.D. conceived the project, supervised all aspects of the work, and edited the manuscript.

## Acknowledgements

D425 and D458 cell lines were generously provided by Dr. Darrel D. Bigner (Duke University Medical Center, NC). SU-DIPG4 was provided by Michelle Monje (Stanford University), and HSJD-DIPG-007 by Angel Montero Carcaboso (Sant Joan de Déu). We likewise thank the University of Colorado Functional Genomics Facility, Genomics Shared Resource, Research Histology Shared Resource, Bioinformatics and Biostatistics Shared Resource, and Animal Imaging Shared Resource for their contributions to this work.

## Ethics Approval

No human subject research was conducted for this study. For vertebrate animal research, University of Colorado Institutional Animal Care and Use Committee (IACUC) approval was obtained and maintained throughout the conduct of the study.

## Data Availability

The accession number for the raw and processed data reported in this paper is GEO: GSE344878.

